# mRNA-Protein Coordination is Contextualized by Metastatic Biological Phenotypes

**DOI:** 10.1101/2025.02.15.638428

**Authors:** Ronit Sharma, Nikolaos Meimetis, Arjana Begzati, Shashwat Depali Nagar, Benjamin Kellman, Hratch M. Baghdassarian

## Abstract

A central goal of conducting omics measurements is to understand how molecular features inform higher-order cell- and tissue-level phenotypes. In particular, multi-omics offers insights into how information encoded by the genome is coordinated through biological layers, resulting in functional outputs^1^. Due to myriad post-transcriptional regulatory processes, the coordination between mRNA and protein cannot be simply reduced to gene-wise correlation. Yet, both modalities have been shown to serve as representations of biological state, and multi-omics integration has been used to improve these representations. Multi-omics approaches typically do not focus on how mRNA and protein features coordinate, but rather use the additional information for improved prediction or feature selection. Here, instead, we showed that standard linear machine learning models provide an understanding of transcriptomic and proteomic coordination in the context of a biological phenotype of interest, in this case cancer metastasis. We find that, in the context of metastasis, a select subset of proteomic features—reflecting a more concentrated signal relative to the broadly distributed transcriptomic signal—offers additional information to that encoded by transcriptomics, as demonstrated by improved model performance when integrating the two modalities and the relative feature importance of proteomics. Top features show a depletion of gene-product overlap across modalities, indicating that the model primarily leverages instances in which the two modalities are providing complementary information with respect to phenotype. However, in instances when both modalities are selected for a given gene product, there is high information consistency that synergistically bolsters phenotype prediction. Altogether, by using model fits that relate both modalities to phenotype, we observe a nuanced coordination of protein and mRNA, in which both modalities tend to provide consistent information about phenotype, yet benefits remain to incorporating a combination of both complementary and reinforcing signals across modalities.

## Introduction

A fundamental goal of systems biology is to elucidate how the coupling of biological layers dictates the flow of information from genotype to phenotype^2^. Pertinently, the extent to which mRNA and protein abundance correlate has been extensively explored, often yielding moderate correlations (∼0.3-0.7)^3,4^. The extent of this concordance, or lack thereof, provides mechanistic insights to myriad post-transcriptional, translational, and post-translational regulatory processes influencing gene product abundance^5,6^. For example, various studies have demonstrated a substantial contribution of protein translation^7^ and degradation^8^ rates in dictating the remaining variance of protein abundance, with such rates combining in specific manners due to synthesis cost considerations^9^. Yet, how mRNA and protein are related with respect to a specific higher-order phenotype of interest has not been extensively explored. Recent proteogenomic efforts, such as pan-cancer analyses of matched transcriptomic and proteomic data, have examined mRNA–protein correlations in phenotypic contexts. However, these analyses largely remain descriptive^10,11^, focusing on correlations independently of the response variable; one study, while associating mRNA-protein concordance with phenotype labels^12^, does not explain how this concordance may be leveraged by predictive models. Yet, interpretable multi-omic predictive models provide opportunity to probe how features contribute within the context of the model fit^13^. Here, we quantitatively associate mRNA–protein coupling with phenotype through predictive models that explicitly leverage both layers.

In parallel, many computational approaches integrate multi-omic measurements. Such approaches tend to emphasize overall predictive gain^14^. Analogous to the lack of phenotypic contextualization in studies of protein-mRNA correlation, multi-omic integration methods that employ an unsupervised approach do not incorporate phenotype in their model fit, instead comparing to phenotypic labels of interest *post-hoc*. While supervised approaches identify shared discriminative features, most of them do not dissect what the model fit reveals about how distinct modalities carry either similar *or* complementary information, or how that information is coordinately leveraged by the model to quantify phenotype. Instead, they focus on selecting features optimized for specific tasks, such as those that maximize correlation across modalities^15^ or interact non-additively^16^ to amplify phenotype. Thus, integrative multi-omic modeling approaches tend to focus on predictive performance rather than the aforementioned topic of mRNA-protein correlation. While many multi-omic integration approaches have their own respective utility, here, we explicitly explore both the correlative and complementary relationships between mRNA and protein features with respect to biological phenotype. To do so, we apply concatenation-based integration^14^ to fit models that simultaneously link proteomic and transcriptomic features to phenotype. Rather than focusing on a specific modeling approach or improving predictive power, we use model fits and resultant predictive power as a proxy for the information each modality carries regarding phenotype.

Using cancer metastasis as a case example of phenotype, we leverage publically available transcriptomic, proteomic, and phenotypic data from DepMap^17,18^ and MetMap^19^, which spans 21 types of solid tumors, to study such coordination. Metastasis causes extensive gene expression regulatory changes, resulting in variable concordance between mRNA and protein. For example, proteogenomic analyses of tumors have shown that genes associated with extracellular matrix remodeling change substantially at the protein level without proportional changes in transcript abundance, reflecting post-transcriptional regulation during metastatic progression^20,21^. Such discordance can arise from altered translation efficiency, microRNA-mediated repression, and ubiquitin–proteasome–dependent protein turnover, processes that are frequently rewired in metastatic cancer cells and contribute to phenotypes such as epithelial–mesenchymal transition (EMT), enhanced motility, and adaptation to new tissue microenvironments^22^.

We test a number of standard linear and non-linear machine learning methods, demonstrating that non-linear models do not outperform linear ones for these sample sizes, lending themselves to simpler interpretability. We identify the best model fit, support vector regression with a linear kernel, and use pathway enrichment analysis to demonstrate that the model captures biologically relevant features. Next, we demonstrate a number of phenotype-contextualized relationships between mRNA and protein both across different molecular species and within the same gene product. Specifically, when assessing each modality independently, we observe that transcriptomics is more informative of cancer metastasis than proteomics. However, our analyses indicate that this difference is not driven by genome coverage or intrinsic modality differences, but instead reflects the specific subset of samples available for proteomic measurements, suggesting that when evaluated on comparable samples the two modalities encode largely similar predictive information. In line with this, model fits and feature selection of individual modalities indicate high similarity. Despite this, surprisingly, we observed that combining the two modalities improves model performance. Overall, correlation alone does not necessarily reflect how much phenotypic information each modality carries or whether combining them yields additional predictive value.

Crucially, the joint modality model fit reveals how proteomics and transcriptomics combine to increase information on metastasis. Improvements from integrating the two modalities arise from combining broadly distributed transcriptomic signals with a smaller set of concentrated proteomic features that contribute largely non-redundant information. At the same time, when both modalities are retained for the same gene product, strong cross-modality consistency reflects synergistic reinforcement of particularly informative phenotypic signals rather than simple redundancy based on high correlation. Altogether, our results indicate that, all else equal, neither protein nor mRNA alone confers inherently greater phenotypic information, but that this information is distributed differently across the genome. Thus, although the modalities often appear redundant when assessed by correlation alone—suggesting that a single modality may suffice for mechanistic insight— the model improves prediction by leveraging a combination of complementary signals across gene products and reinforced signals where mRNA and protein are strongly consistent.

## Results

### Linear Models Perform At Least as Well as Non-Linear Models in Predicting Metastatic Potential from Transcriptomics

In order to determine whether global gene product abundances are informative of metastasis, we leveraged DepMap’s cell line measurements^17^. Specifically, we fit a dataset containing 481 samples on a set of machine learning models that take as input protein-coding genes measured by RNA-sequencing and proteomics and predict as output metastatic potential reported by MetMap 500^19^. Given our goal of understanding features in relation to each other, we first set out to ask whether more easily interpretable linear models could offer comparable performance to more complex non-linear models in datasets of this size; this is particularly relevant as most datasets with transcriptomics, proteomics, and phenotypic measurements do not exceed this size. Given its availability, we first focused on transcriptomics to see how well that modality alone could recover phenotypic information.

After hyperparameter tuning and rank-ordering of 9 linear and non-linear machine learning models by median performance (Fig. S1a, see Methods for Details), we further assessed the top three models (support vector regression (SVR) with radial basis function, polynomial, and linear kernel) using consensus hyperparameters, alongside single and ensemble neural nets. Despite potentially complex relationships between molecular features and cell phenotypes such as cancer cell metastasis, we observed no significant differences between the linear SVR and the non-linear models (Mann-Whitney U, Benjamini-Hochberg FDR correction, q ≥ 0.69, Cohen’s d ≤ 0.22), including neural nets (Fig. 1a, Fig. S2a). These results are further supported by the lack of significant differences (q ≥ 0.86) between all 9 model predictions during hyperparameter tuning using nested cross-fold validation (CV) (Fig, S1a, Table 3).

**Fig. 1:**
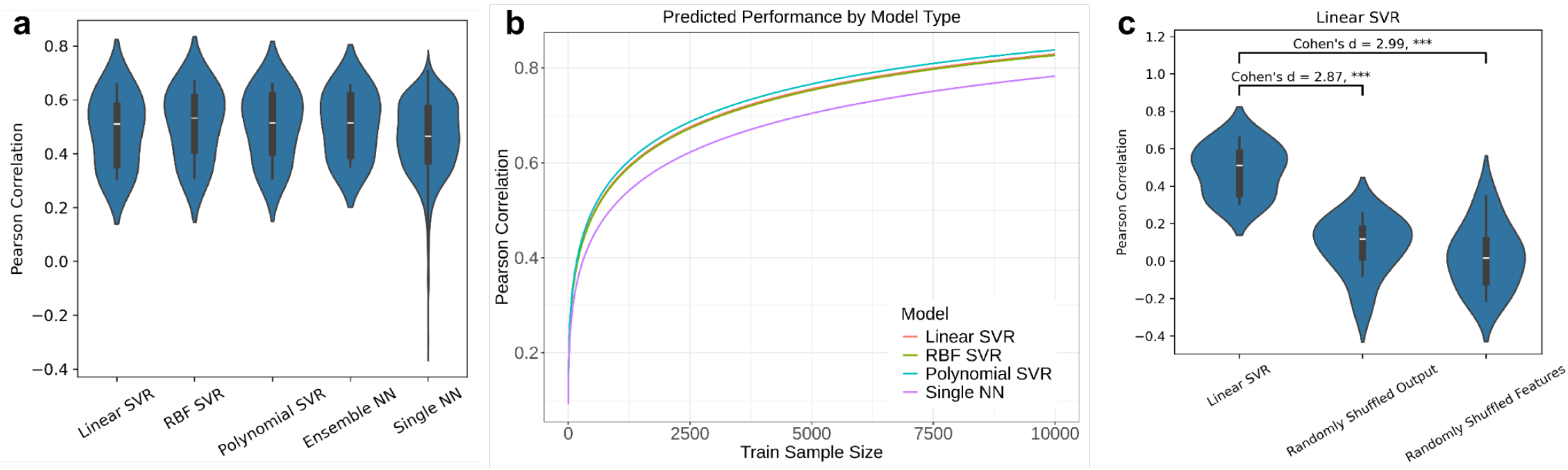
Linear models exhibit predictive power for metastasis from transcriptomics. All statistical significance is determined using a Mann-Whitney U (MWU) test and the FDR is controlled for multiple testing using the Benjamini-Hochberg correction (**** q≤10^-4^, *** q≤10^-3^, ** q≤0.01, * q≤0.1). Comparisons that are not significant are not annotated. Models’ predictive performance is assessed by Pearson correlation. Violin plots visualize performance distribution output from 10-fold CV. **a** Comparison of consensus linear SVR model to other consensus SVR models’ and neural nets’ (x-axis) performance (y-axis) (two-sided MWU test). No comparisons have q ≤ 0.1. **b** Estimates of Pearson correlation (y-axis) for each consensus model type across training sample size (x-axis) from 1 to 10,000 training samples. Estimates are derived from a power function regression modeling Fisher z-transformed Pearson correlation as a function of Train Sample Size and including model type as a covariate. **c** Consensus Linear SVR model performance (y-axis) compared to random baselines (x-axis) (one-sided MWU test).

Because the number of features used is much larger than the number of available samples, and because more complex models with many parameters such as neural nets require large sample sizes, we conducted a power analysis to determine how each model’s performance changes with sample size (Fig. S4, see Methods for details). First, we determined that the change in performance across sample size is best explained by a power function, as opposed to a linear or exponential function (Table 1). Next, we fit a power function regressing model performance against the training sample size, and we included the model type as a covariate to assess differences between them (Fig. S5). When controlling for sample size, this regression estimates the nonlinear SVRs have a slightly (10^0.01^ = 1.02-fold) higher Fisher z-transformed Pearson correlation than the linear SVR. However, practically speaking, this difference is negligible (Fig. 1b). For example, when extrapolating to 10,000 samples, all SVRs are estimated to have a Pearson correlation of approximately 0.83-0.84. Furthermore, the single neural nets are estimated to perform more poorly than the linear SVR; the Fisher z-transformed Pearson correlation is 0.9 (10^-.05^) times lower than the linear SVR, predicting a Pearson correlation of 0.78 at 10,000 samples. Results for mean squared error (MSE) are qualitatively similar (Fig. S5). This indicates that the sample size is a much more substantial limiting factor in model performance than model type.

Since SVR models were the top performers (Fig. S1a), and because non-linear kernels did not outperform the more interpretable linear kernel, we proceed with downstream analyses using the linear SVR. With a mean 10-fold CV Pearson correlation of 0.51, we confirm that the consensus linear SVR model outperforms random baselines in which the output values or input features were randomly permuted prior to fitting the model with the same hyperparameters (Fig. 1c, Fig. S2b). Altogether, these results indicate that machine learning models can explain ∼25.5% of the variance (quantitated by the median 10-fold CV test predictions’ coefficient of variance of the consensus linear SVR) in metastatic potential from transcriptomics, and that, for these data, a more interpretable linear model suffices in capturing this relationship.

### Transcriptomics and Proteomics Encode Largely Overlapping Information but Yield Improved Predictive Performance When Combined

Next, because protein-level gene abundance may contain additional information about cell phenotypes independent of that available in RNA-level gene abundance^23^, we asked how models perform on proteomics data. We again identified consensus models for the top 3 performers using proteomics (Fig. S1b). Because the proteomics data had fewer samples, we compared it to the transcriptomics consensus linear SVR model run on an equal sample size from the power analysis. We found that the transcriptomics model outperformed all top three proteomics models as measured by MSE (Fig. 2a, Fig. S3a, Fig. S6). Additionally, proteomics models did not outperform the transcriptomics one as measured by Pearson correlation. We note that, as in the transcriptomics fit models, those fit on either proteomics or both modalities also indicated that linear SVRs were the top-ranked linear models and not outperformed by non-linear models (Fig. S8); consequently, for ease of interpretability and more direct comparisons, we proceed with downstream analyses using consensus linear SVRs.

**Fig. 2:**
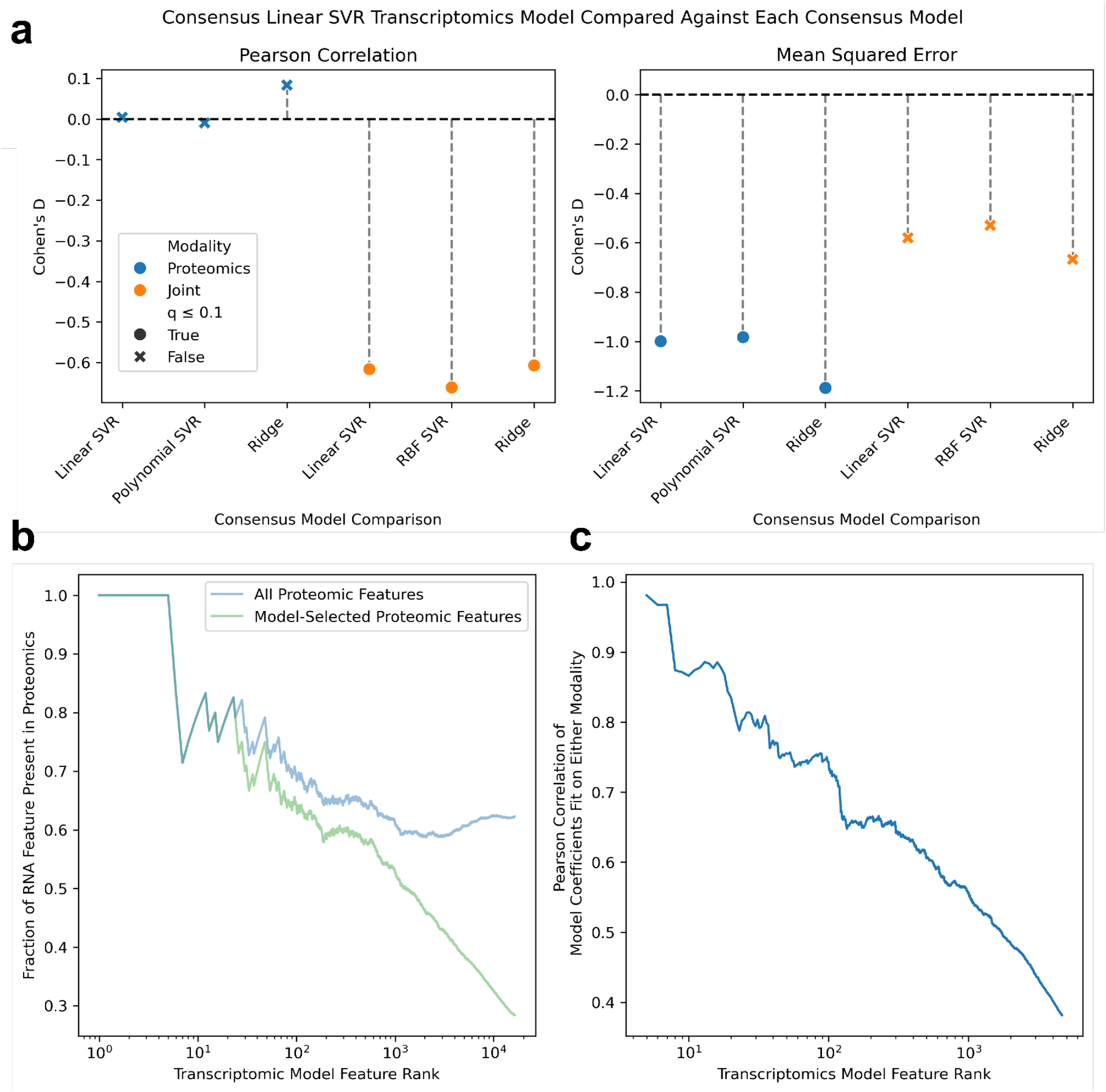
Proteomics and Transcriptomics Offer Consistent Views of Metastasis. **a** Lollipop plots visualize comparisons between top consensus models (x-axis) identified in proteomics (blue) or joint omics (orange) with respect to the consensus transcriptomic linear SVR. Models’ predictive performance is assessed by Pearson correlation (left panel) and mean squared error (right panel). Comparison effect sizes are quantified using Cohen’s d (negative values mean the metric effect was smaller in the transcriptomic model and positive values mean the metric effect was larger in the transcriptomic model). Significance (two-sided Mann-Whitney U test, Benjamini-Hochberg FDR correction, q ≤ 0.1) is indicated by the scatter point shape. Performance for each consensus model (x-axis) is assessed by 10-fold CV, and compared against the linear SVR performance at the same sample size using the power analysis previously described. **b** Line Plot of the fraction of transcriptomic features shared with proteomics (y-axis) as a function of the number of transcriptomic features at a given rank-order (x-axis). Feature ranks are obtained from fitting the linear SVR model with consensus hyperparameters on the entire transcriptomic dataset and ordering by absolute value of the model coefficient. Coloring displays all proteomic features (blue) and the top 5,000 from feature selection in the consensus linear SVR (green). **c** Line plot of the Pearson correlation at a given transcriptomic feature rank between transcriptomics and proteomics consensus linear SVR coefficients. Features are subsetted to those shared between the two models (4,654 shared features between 5,000 proteomic and 16,371 transcriptomic features), and the x-axis displays the relative transcriptomic feature rank.

Because proteins are often viewed as more proximal to phenotype than transcripts, we asked whether transcriptomics may outperform proteomics due to its more comprehensive genome-wide coverage (16,371 transcriptomic features compared to 10,969 proteomic features). We first quantified the extent of feature overlap between the two modalities. If reduced genome coverage were the primary limitation of proteomics, we would expect informative transcriptomic features to be increasingly absent from proteomics at higher feature ranks. Instead, we observed substantial overlap at high ranks: 69.0% and 64.4% of proteomic features were present among the top 100 and 500 transcriptomic features ranked by the transcriptomic model, respectively (Fig. 2b, blue line). Moreover, when highly ranked transcriptomic features were available in the proteomics dataset, they were frequently selected by the proteomics model; 64.0% and 58.2% of selected proteomic features were present among the top 100 and 500 transcriptomic features, respectively (Fig. 2b, green line).

This suggests that, for shared high-importance features, the two modalities capture similar predictive information. Consistent with this, model coefficients for overlapping features showed strong agreement, with a Pearson correlation of 0.74 among the top 100 shared features ranked by the transcriptomic model (Fig. 2c). Together, these results indicate that features most informative to transcriptomics are largely available to proteomics and are utilized similarly by both models, arguing against genome coverage alone as the primary driver of performance differences.

However, given that there was still a substantial portion of top transcriptomic features missing in proteomics, we more explicitly assessed this using a 10-fold CV of the intersecting features. We found that, when the comparison is restricted to the intersecting set of samples between transcriptomics and proteomics (247 intersecting samples between 248 proteomic and and 481 transcriptomic samples), performance differences between modalities are lost, regardless of whether models are trained on all available features or only their intersection (Fig. S9a-b). Moreover, within this intersecting sample set, performance is comparable between models fit on all versus intersecting features within each modality. (Fig. S9c-d, see Supplementary Results - “Transcriptomic and Proteomic Comparison Intersecting Features and Subsets” for an extended discussion). Altogether, these results indicate that differences in feature availability could not account for the observed performance gap. Instead, performance seems to be constrained by the specific subset of samples available to proteomics, regardless of modality or feature availability.

Next, we asked whether a joint model incorporating both modalities could leverage complementary information, despite the substantial agreement observed between single-modality models. Surprisingly, despite the fact that (i) the subset of samples available to proteomics is a major factor of reduced model performance independent of sample size, modality, and feature availability and (ii) shared features were utilized similarly by single-modality models, we found that the top consensus joint omics models consistently outperformed the consensus transcriptomic model by Pearson correlation (medium to large effect sizes, -0.61 ≥ Cohen’s d ≥ - 0.66; Fig. 2a, Fig. S3b, Fig. S1c). Notably, joint multi-omics models did not outperform transcriptomics when evaluated by MSE; although differences were not statistically significant, effect sizes indicate slightly worse MSE performance for the joint models relative to transcriptomics (−0.58 ≥ Cohen’s d ≥ −0.67; Fig. 2a). Despite this, median Pearson correlations for the joint multi-omics model evaluated on the 247 shared samples were approximately 0.5—comparable to those achieved by transcriptomic models trained on the full set of 481 samples. In contrast, transcriptomic models evaluated under the power analysis at matched sample size achieved a median correlation of 0.43, while proteomic models trained on 248 samples achieved a median correlation of 0.42. Together, these results indicate a substantial quantitative improvement in predictive performance when incorporating both modalities, corresponding to an increase of approximately 6–7 percentage points in variance explained.

All consensus joint omics models performed better than random baselines (Fig. S7). Altogether, these findings suggest that integrating multiple modalities can yield additional phenotypic information, even when the modalities themselves exhibit substantial overlap, a theme we explore further in the following section.

### Multi-omic Modeling Captures Metastasis-Associated Coordination Between Modalities

Given the improvement in Pearson correlation achieved by joint multi-omics models, despite substantial concordance in feature selection and model fit across modalities, we next asked how the two modalities are jointly leveraged by the model—through discrepancies or reinforcement of shared information. A joint model incorporating both modalities can provide insight into how mRNA–protein relationships are coordinated with respect to phenotype.

We first checked whether the model fit is capturing features relevant to metastatic biology. To do so, we rank-ordered features by the absolute value of their coefficient in the model fit on the entire dataset and ran an over-representation analysis (ORA) of the top 500 features. Many of the overrepresented pathways are related to metastasis. For example, “positive regulation of locomotion” and “chemotaxis” are explicitly related to cell motility and directional migration. Cell–cell adhesion and integrin-mediated adhesion are key processes in metastasis, as they regulate how cancer cells detach from the primary tumor, interact with the extracellular matrix, and migrate to invade and colonize distant tissues. Finally, many terms are associated with ECM remodeling, which enables tumor cells to degrade and reorganize the surrounding matrix, facilitating migration.

Next, we asked how the proteomic and transcriptomic data modalities were combined with regards to the model fit. Much like feature selection for the individual modalities, hyperparameter tuning consistently selected more transcriptomic features than proteomic ones, resulting in a consensus model with all 16,371 transcriptomic features selected but only 1,475 proteomic features, representing only 8.3% of all features incorporated in the model. The selection of only a small subset of proteomic features but all transcriptomics features was consistent across model types (Table 2). However, proteomic features represent 36.60% of the top 500 features (odds ratio 7.17; Fisher’s exact test one-sided p-value 6.96e-7; Fig. 3b). These results indicate that while proteomics comprise only a small subset of the features, they represent important additional information for the model. This aligns with our previous observations regarding the boost in performance provided by using the joint omics data (Fig. 2a, Fig. S3b) and the lower effective dimensionality of proteomics as compared to transcriptomics given feature availability (Fig. 2c; Supplementary Results - Transcriptomic and Proteomics Comparison Intersecting Features and Subsets). Given the variance of within-gene correlation between mRNA and protein abundance^3,23,24^, we also asked the extent to which gene products overlapped between these two modalities in the context of the joint omics model fit. While there was nearly complete overlap across all selected features, only 26.1% of proteomic features overlapped with transcriptomic features in the top 500, indicating a significant depletion (odds ratio 0.05, Fisher’s exact test one-sided p-value 8.95e-68) of gene product overlap between mRNA and protein (Fig. 3c). This depletion indicates that the joint model preferentially selects non-redundant gene products across modalities.

**Fig. 3:**
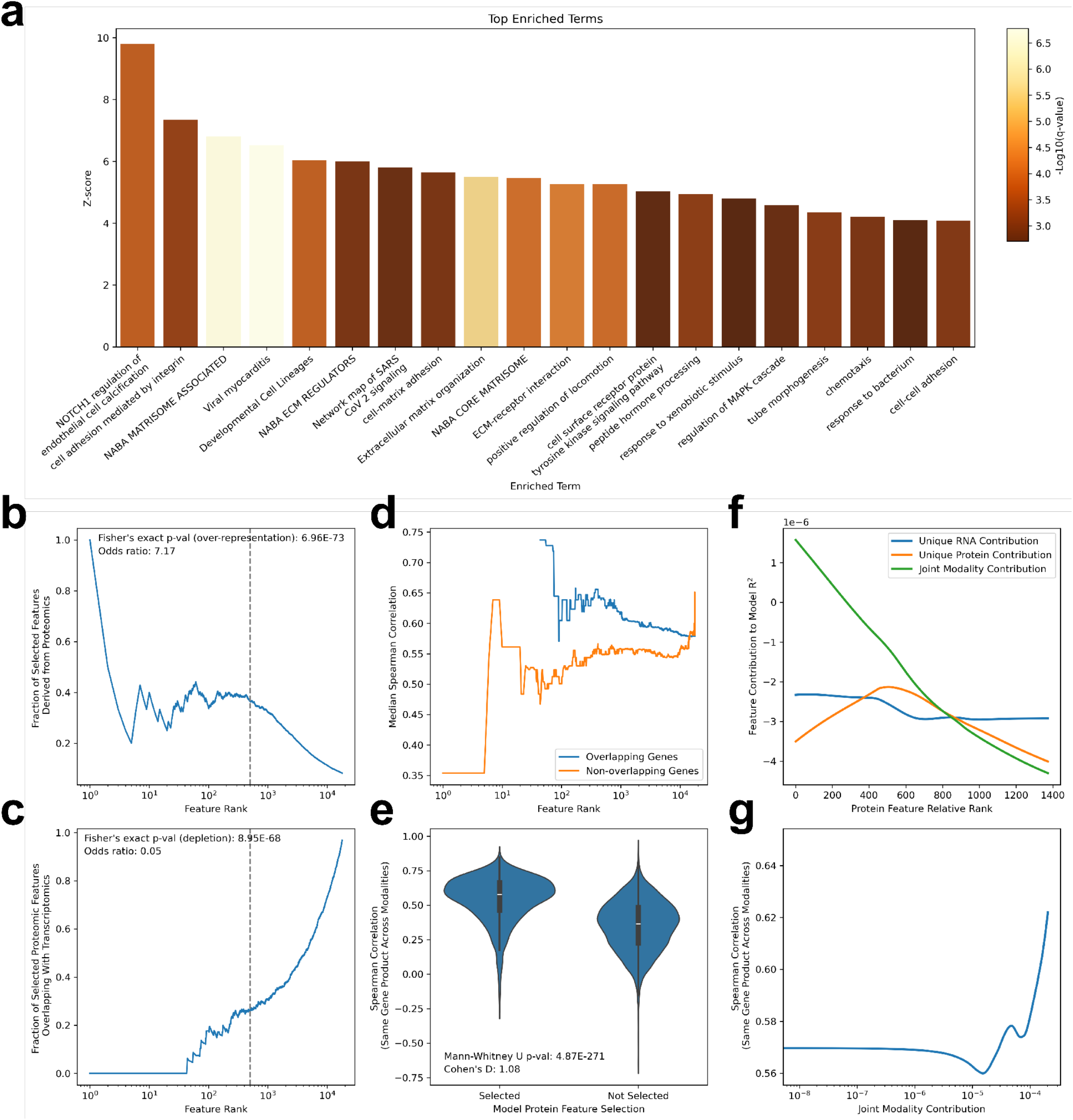
Proteomics and transcriptomic features complement each other in driving metastatic programs. **a** Barplots of the top 20 Metascape over-represented summary terms for highly ranked features in the consensus linear SVR. Terms are rank-ordered by z-score and colored by -log_10_(q-value). Z-scores are aggregated by their average across individual members of each summary term. **b** Line Plot of the fraction of selected features derived from proteomics (y-axis) as a function of the number of features at a given rank-order (x-axis). Features are rank-ordered by the absolute value of the model coefficient. The gray dashed line represents a rank of 500, at which statistics were calculated. **c** Line Plot of the fraction of selected proteomic features (y-axis) that overlap with selected transcriptomic ones as a function of the number of features at a given rank-order (x-axis). For a proteomic feature at a given feature rank, we define a corresponding overlapping transcriptomic feature as one that has a shared gene product and a feature rank at or below (“higher rank” - more important) that of the proteomic feature. Conversely, a non-overlapping transcriptomic feature is one that has a shared gene product with the proteomic feature and a feature rank above (“lower rank” - less important) that of the proteomic feature. **d** Line Plot comparing differences in proteomic-to-transcriptomic feature correlation for overlapping or non-overlapping genes (y-axis) at a given rank-order (x-axis). r_overlapping_ is calculated by identifying all proteomic features at a given rank that have overlapping (shared gene-product) transcriptomic features at or below that rank, calculating the Spearman correlation between each feature pair across samples, and taking the median across feature pairs. r_non-overlapping_ is similarly calculated, but instead identifies all proteomic features at a given rank whose corresponding transcriptomic features are above that rank. In instances of many-to-one ID mappings between modalities, only the highest ranking feature within a modality was retained for this analysis. To avoid spurious correlations due to low-abundance, noisy features, we set global per-modality cutoffs: for each features in a modality, we calculated the 10th percentile abundance value across samples, took the median across features, and, in each feature pair for correlation, excluded samples if either modality measurement fell below this value. **e** Violin plots compare Spearman correlations for proteomic features selected or excluded by the model. Correlations were calculated as in (d), taking the sample-wise Spearman correlation of features with shared gene products between the two modalities. **f** Line Plots show the change in coefficient of determination relative to the full model following feature removal at a given proteomic feature rank: removal of both a proteomic feature and its corresponding transcriptomic feature sharing the same gene product (green), removal of the proteomic feature alone (orange), or removal of the corresponding transcriptomic feature alone (blue). Larger magnitudes indicate a stronger influence of the feature (or feature pair) on model performance, with positive values reflecting a helpful contribution to prediction and negative values indicating detrimental effects. **g** Line Plot shows the relationship between the cross-modality Spearman correlation (as calculated in panel d) and the joint modality contribution (as calculated in panel f) for proteomic features and their corresponding transcriptomic feature.

This preferential selection of one modality over another suggests that mRNA and protein measurements often encode overlapping phenotypic information, consistent with the strong agreement observed between single-modality model fits (Fig. 2c). Consequently, we set out to assess the extent to which complementarity between mRNA and protein information is accounted for by the model fit. In instances where a proteomic feature was selected over its corresponding transcriptomic feature sharing the same gene product at a given feature rank (i.e., *non-overlapping*), the two measurements may encode *consistent* phenotypic information, with the model preferentially selecting one representative. In such cases, selection of the proteomic feature may reflect differences in signal quality or relevance to metastatic phenotype relative to its transcriptomic counterpart.

Conversely, when both the proteomic feature and its corresponding transcriptomic feature sharing the same gene product were selected at a given feature rank (i.e., *overlapping*), this suggests that the two modalities may contribute *complementary* information for that gene product.^23^ To explicitly test this, we assessed how the Spearman correlation between transcriptomic and proteomic features differed between overlapping and non-overlapping cases (Fig. 3d), using higher correlation as a proxy for greater consistency and lower correlation as a proxy for greater complementarity. Overall, feature pairs exhibited high correlations in both cases.

Surprisingly, at a given feature rank, overlapping features tended to show higher Spearman correlation than non-overlapping ones. In conjunction with the observed depletion of gene product overlap (Fig. 3c), these results indicate that the joint model typically resolves highly consistent mRNA–protein signals by preferentially selecting a single modality. However, in cases where both modalities are retained for a given gene product, the corresponding features exhibit particularly high cross-modality consistency relative to cases of preferential selection. This suggests that joint inclusion reflects reinforcement of an especially informative phenotypic signal, with both modalities contributing concordant information that the model deems valuable to retain.

Extending this analysis to compare proteomic features that were selected or excluded yielded similar results: selected proteomic features exhibit significantly higher consistency with their transcriptomic counterparts than those that were excluded (Fig. 3e).

In the context of multi-modal analysis with a phenotypic response variable, these results indicate that feature selection need not prioritize divergent signals across modalities. Given the unexpected result that model fit prioritized consistent features in instances where gene products overlapped, we next further probed the contribution of such features to model performance using a dominance analysis that decomposed the predictive contributions of each modality and their combination at a given feature rank. Specifically, for selected proteomic features with corresponding transcriptomic features sharing the same gene product, we systematically removed the proteomic feature, its corresponding transcriptomic feature, or both, re-fit the model, and quantified the resulting change in the coefficient of determination (Fig. 3f). Overall, each individual feature contributes only a small amount to the variance explained in metastatic phenotypes, indicating that predictive performance arises from aggregating information across many features. At higher ranks, the combined contribution of transcriptomic and proteomic features tends to exceed that of either modality alone, such that even when an individual modality is weakly informative or detrimental under refitting, the joint contribution remains beneficial. Notably, the joint contribution decreases with increasing protein-feature rank, indicating that higher-ranked protein features more frequently form cross-modality pairs that contribute strongly to metastatic phenotype prediction, even when the information encoded across modalities is largely consistent (Fig. 3d). In fact, we observed a positive association between cross-modality consistency and the joint contribution of both modalities (Fig. 3g), further supporting the notion that, when the model prioritizes overlapping features, strong cross-modal coherence encodes biologically meaningful variation relevant to metastatic potential. Thus, correlation alone does not fully capture phenotypic information content, and high cross-modality correlation does not preclude substantial marginal benefit from integrating a second modality.

In summary, Fig. 3b–c together suggest that the primary source of improvement in the joint omics model arises from combining broadly distributed transcriptomic information with a small set of concentrated, largely non-redundant proteomic features. Yet, Fig.3d-g indicates that in cases where both modalities are retained for the same gene product, high cross-modality consistency rather than complementarity reflects reinforcement of an especially informative phenotypic signal.

## Discussion

In this study, we used metastatic potential as a lens to examine how transcriptomic and proteomic layers encode phenotype and how they coordinate when leveraged by predictive models simultaneously. First, we demonstrate that linear machine learning models can use cancer metastasis as an example phenotype to reveal feature information contextualized by a phenotype as well as more complex non-linear models, including ANNs. However, this speaks in part to the large variance and moderate predictive power of these data, yielding what appears to be an upper limit to the information standard machine learning approaches can extract.

Next, we observed that transcriptomics predictions tend to outperform proteomic ones. When using the same starting set of genes, proteomics did not outperform transcriptomics, despite proteins typically being considered more proximal to mechanism or phenotype than mRNA. Instead, performance differences were largely attributable to the specific subsamples available to either modality, rather than feature availability; consequently, at the same sample size, models performed similarly fit on either modality regardless of whether the intersection of features was taken. Notably, the proteomics pipeline typically selected substantially fewer features, consistent with a more concentrated signal relative to the broadly distributed signal observed in transcriptomics. Additionally, each sequencing technology has distinct limitations; consequently, other contributing factors may include limits-of-detection preventing quantification of low-abundance proteins, which is not as much of an issue in RNA-seq^25^ and higher noise in proteomics measurements^26^. We note that our observations regarding discrepancies or similarities in model performance between proteomic and transcriptomic modalities may not necessarily generalize to other datasets or phenotypes.

Combining proteomics and transcriptomics offered an increase in predictive performance as measured by Pearson correlation. The most informative features in our linear joint omics model were associated with processes relevant to metastasis such as cell migration and chemotaxis, indicating that despite moderate predictive performance the model captures biologically meaningful relationships between metastasis and the molecular profiles of the cell lines. The model fits tended to combine features from both modalities in a largely non-redundant manner: there was significant enrichment of proteins among the top 500 features together with a significant depletion in modality overlap of gene products. This suggests that a subset of proteomic features provides information not captured by their transcriptomic counterparts. Consistent with this, dominance analysis indicated that higher-ranked cross-modality feature pairs contributed disproportionately to predictive performance, even when individual features contributed only modestly to the variance explained. Interestingly, in cases where both modalities were retained for the same gene product, high cross-modality consistency was associated with stronger joint contributions to prediction, suggesting that concordant mRNA–protein signals can reinforce particularly informative phenotypic variation rather than representing simple redundancy. Thus, across cell lines, the model fit is accounting for gene-specific variance in correlation^3^ between mRNA and protein, where both complementary signals across gene products and reinforced signals within gene products contribute to metastasis prediction. The presence of non-linear relationships and network effects^27^ between mRNA and protein may further explain instances in which the two modalities are distinctly informative^23^ of metastasis.

These results are based on omics measurements of cancer cell lines, which might have substantial differences from *in vivo* tumors. *In vivo* metastasized tumors contain heterogeneous microenvironments influenced by mutational burden, interactions with tissue-specific cell types and the immune system, and nutrient availability and hypoxia^2,28^. Similarly, the metastatic potential was quantified in mice, and therefore might not directly translate to human metastatic phenotypes due to factors such as functional differences in orthologs^29^, distinct global responses to perturbation^30^, and pathogenesis through disease mechanisms that are subdominant in one species^31^. While molecular processes in these simpler models do not fully reflect *in vivo* responses, our pathway enrichment results indicate that the model still captures biologically relevant features. Nonetheless, tools^30,32^ that can contextualize and translate the omics profiles of cancer preclinical models to clinical responses can help improve predictive power, interpretability, and clinical relevance. Similarly, while our model is tissue-agnostic, explicitly accounting for the tissue-specific metastatic potential available in MetMap may improve specificity and predictive power. Altogether, these findings highlight the value of multi-omics data integration in interpretable models, not only to enhance predictive accuracy but also to reveal how features from different modalities jointly associate with a phenotype of interest. With the advent of high throughput proteomic platforms measuring thousands of proteins^33,34^, multi-omic measurements with much broader coverage are emerging. This enables researchers to use multi-omic machine learning models to uncover the complementarity and level of interaction of each modality more thoroughly, generating better mechanistic hypotheses about phenotypes of interest.

## Methods

### Transcriptomic and Proteomic Data Processing

Metastatic potential for 488 samples reported by MetMap 500^19^ was downloaded from https://depmap.org/metmap/data/index.html.

Batch-corrected RNA-sequencing data of protein-coding genes for the DepMap^17^ cell lines was downloaded in log-normalized transcripts per million (https://depmap.org/portal/data_page/?tab=allData&releasename=DepMap%20Public%2024Q4&filename=OmicsExpressionProteinCodingGenesTPMLogp1BatchCorrected.csv). 481 samples overlapped out of those reported in MetMap 500 and the 1673 reported in the RNA-sequencing dataset. 2,767 of 16,371 low-abundance features were filtered for mean values < 0.1.

Normalized protein expression data reported in Table S2 of Nusinow et al.^18^ were used. Proteins with missing values in more than 80% of samples (1,786 of 12,755 features) were removed. The remaining missing values were imputed in Perseus (version 2.1.3.0) as follows: the protein distribution was transformed by 1) subtracting 1.8 standard deviations from all values and then 2) multiplying all values by 0.3 to shrink the standard deviation; next, imputed values were drawn by random sampling from this transformed distribution. Three cell lines had two samples originating from different ten-plexes, which were dropped. Finally, 248 samples overlapped out of those reported in MetMap 500 and the 372 reported in the proteomics dataset. 247 samples were present in both the transcriptomics and proteomics data.

### Model Fitting and Hyperparameter Tuning

The following models were each fit on the data, with omics as the input and metastatic potential as the output: partial least squared (PLS) regression, Ridge, Lasso, ElasticNet, Support Vector Machine (SVM) with a linear kernel. Similarly, the following nonlinear regression models were fit on the data: SVM with polynomial and RBF kernels, k-Nearest Neighbors (KNN), and Random Forest.

Models were either fit on transcriptomics separately, proteomics separately, or the two datasets jointly. The following preprocessing steps were always conducted on either modality separately: the top features were selected according to those that had the highest mean-variance residual. Specifically, features are ranked by the residuals from a LOWESS-fitted mean–variance trend, identifying those whose variance is higher than expected given their mean expression. Feature selection is only conducted on training data and proceeded by mean-centering. The number of features used by the model was one of the hyperparameters explored (see Table 2 for a comprehensive accounting of hyperparameters explored).

Each model type was fit on the same 10 train-test splits using 10-fold CV. Within each fold, an inner 5-fold CV on the training data was used to identify the best set of hyperparameters (Table 2, sheet 1). Hyperparameter tuning was conducted using Optuna^35^ with 100 trials to minimize the average mean squared error. The “SuccessiveHalvingPruner” pruner was used alongside a custom sampler. The custom sampler used the CmaEsSampler on hyperparameters when possible, and the TPESampler otherwise. To prevent convergence to a local minimum, every 20 trials, hyperparameters were also randomly sampled using the TPESampler.

Upon identification of the best set of hyperparameters, model predictions were assessed on the outer fold test data using Pearson correlation and mean squared error (Fig. S1). This resulted in a unique set of hyperparameters for each fold for each model type (Table 2, sheets 2-4). We rank-ordered each model as follows: we aggregated performance across the outer 10 folds by the median test MSE and test Pearson correlation, rank-ordered models by each respective metric, and calculated a consensus rank as the median across the two metrics. In cases where models shared the same consensus rank, ties were resolved using additional consensus ranks derived sequentially from the standard deviation and then the mean across folds (i.e., if the standard-deviation consensus also tied, the mean-based consensus was used). A single consensus model for each of the top three performing model types for both performance metrics was generated as follows: 1) for discrete hyperparameters, the mode across 10-folds was used, 2) for continuous hyperparameters, the mean value across the 10-folds was used, and 3) for integer hyperparameters, when a majority of observations shared the same value (defined as the most frequent value occurring in ≥50% of samples), we assigned the mode. When no such dominant value existed, indicating more uniform distributions across folds, we instead used the mean rounded to the nearest integer. Once a consensus model was identified, it was assessed using the same performance metrics on a different set of folds using 10-fold CV (Fig. 1A, Fig. S2A, and Fig. S3). A 10-fold CV was also used to assess single and ensemble artificial feed-forward neural networks (ANNs) (see Supplementary Methods for details).

### Power Analysis of Transcriptomic Model

To assess how model performance scales with training sample size, we conducted a power analysis using the top-performing consensus models fit on transcriptomics. We trained models on 10–100% of the training data in 10% increments using 10-fold CV, repeating each subsample 100 times to generate stable estimates.

Performance was evaluated using mean squared error (MSE) and Pearson correlation, the latter Fisher z-transformed prior to modeling. We compared linear, power, and exponential fits to describe the relationship between sample size and model performance, selecting the best function based on adjusted R^2^, AIC, and BIC. Power functions best explained the performance trends across nearly all models and metrics. We then fit a power regression model with model type as a covariate to quantify performance differences across sample sizes. For additional details, see the Supplementary Methods.

## Author Contributions

R.S.: Conceptualization, Transcriptomic Analysis, Writing N.M.: Conceptualization, Neural Network and Feature Analysis, Writing, and Editing A.B.: Feature Analysis, Writing S.D.N.: Power Analysis, Writing B.K.: Conceptualization H.M.B.: Conceptualization, Analysis, Writing, Editing, and Supervision.

## Acknowledgements

We would like to thank Luka Karginov and Brian Joughin for helpful discussions regarding correlation networks and feature selection, respectively. We would like to thank Douglas Lauffenburger and Nathan Lewis for helpful discussions regarding the relationships between transcriptomics and proteomics. H.M.B is funded as a MIT-Novo Nordisk Artificial Intelligence Postdoctoral Fellow and is supported by a Cancer Research Institute Immuno-Informatics Postdoctoral Fellowship (CRI12812). N.M. is a 2024-2025 Takeda Fellow.

## Data Availability

All data is publicly available through DepMap (https://depmap.org/portal/) and CCLE (https://sites.broadinstitute.org/ccle/).

## Code Availability

Code for all analyses in this manuscript are available on Github https://github.com/hmbaghdassarian/metastatic_potential.

