## Supplementary Information and Figures for "mRNA-Protein Coordination is Contextualized by Metastatic Biological Phenotypes"

#### Supplementary Methods

##### Neural Network Construction and Training

For transcriptomics, alongside the consensus models, we also fit single and ensemble artificial feed-forward neural networks (ANNs) on the 10-fold CV. Single ANNs consist of one initial dropout layer, with a dropout rate of 0.5, and four blocks which contain in series: a fully-connected linear layer, a batch normalization layer with (momentum=0.2), a non-linear activation function (ELU), and a dropout layer with a dropout rate of 0.25. The final layer is a Gaussian regression layer that predicts the mean ($\bar{y}_i$) and a variance ($y_{\sigma_i}$) of the metastatic potential, rather than a single value. This treats the metastatic potential as a sample from a Gaussian distribution with the predicted mean and variance, and the model is trained by minimizing the negative log-likelihood criterion[1] given by: $-log(p_\theta|X_n) = -\frac{1}{2}log\sigma_\theta{}^2(x) - \frac{1}{2\sigma_\theta{}^2(x)}(y - \mu_\theta(x))^2 + constant$ and minimized through the following loss function: $Loss = \frac{1}{2}logy_{\sigma_i} + \frac{1}{2y_{\sigma_i}}(y_{true} - \underline{y_i})^2 + 10^{-6}$. We note that for the power analysis (see next section for details), we do not use a Gaussian regression layer as the final layer, but rather a simple linear feed-forward neural network layer. Each layer reduces the dimensionality of the data progressively to 1024, 256, 32, and 16 nodes. Models were trained for 100 epochs with a learning rate of 0.001 and a batch size of 40 using the Adam optimizer. To avoid overfitting L2-regularization was applied to the models' parameters with a regularization strength of 0.01.

Next, an ensemble of 20 of these ANNs is trained. The output of this ensemble is also a Gaussian, with mean and variance calculated from the uniformly weighted mixture of each model. The final prediction of the model corresponds to the mean of the mixture: $y_\mu = \frac{1}{N}\sum_{i=1}^{N}\bar{y}_i$ . The variance of the mixture is defined as: $\sigma = \sqrt{\frac{1}{N}\sum_{i=1}^{N}(y_{\sigma_i} + \underline{y_i}^2) - y_\mu{}^2}$ .

##### Power Analysis of Transcriptomic Model

To assess how the top-performing, transcriptomics-fit consensus models are affected by sample size, we conducted a power analysis. We first split data into training and test splits using 10-fold CV. For each fold, we subsetted training data between 10% to 100% of the sample size at intervals of 10%. We also conducted the power analysis at the sample sizes of the proteomics and joint omics datasets (248 and 247 samples respectively). With the exception of the entire training sample size, for each interval, we selected 100 random subsets to fit the model on, resulting in a total of 1000 test predictions per training sample size.

Next, we quantified the relationship between prediction performance, sample size, and model type. First, we determined whether a linear, power, or exponential function best explains the relationship between model performance and sample size. To do so, we fit linear models for each of MSE or Pearson correlation regressed on sample size for each model type individually. Prior to model fitting, for Pearson correlation, we applied Fisher's z-transformation. For power functions, we fit the relationship $m = aN^b$, where $m$ is the model performance (Fisher z-transformed Pearson correlation or MSE), $N$ is the training sample size, and $a$ and $b$ are the parameters for the power function. We log-transform this data to fit a linear regression to it as follows: $\log_{10}(m) = \log_{10}(a) + b*\log(N)$. Similarly, for exponential functions, we fit the relationship $m = a*e^{bN}$. We log-transform this data to fit a linear model to it as follows: $\log_e(m) = \log_e(a) + bN$. Next, we assessed goodness of fit using the adjusted $R^2$, Akaike information criterion (AIC), and Bayesian information criterion (BIC). We observed that on average across the predictive models, power functions best-explained performance by adjusted $R^2$ for both Pearson correlation and MSE, and they also best explained performance by AIC and BIC for MSE (Table 1).

As such, we proceeded with a power function to fit the following regression in order to assess differences in model types' predictive performance across sample size: $\log_{10}(m) = \log_{10}(a) + b*\log(N) + model\_i$, where $i$ is the model type and the Linear SVR is the reference model. When back-transforming to the power function, these fitted parameters can be interpreted as follows: $m = aN^{b}*10^{model\_i}$. For the Linear SVR to be the reference, $model\_linearSVR = 0$, such that $m = aN^{b}*10^{0}$, which simplifies to $m = aN^{b}$. Thus, each other model covariate is a scaling of model performance by $10^{model\_i}$ relative to this reference.

### Supplementary Results

#### Transcriptomic and Proteomic Comparison Intersecting Features and Subsets

The consensus proteomics models tended to have comparable performance to the consensus transcriptomics linear SVR in terms of Pearson correlation (Fig. 2a, S3a). Furthermore, in contrast to the transcriptomics models, which tended to select all 16,371 available features, proteomics models managed to do so with only 5,000 of 10,969 available features selected (Table 2, sheet 5). As such, we hypothesized that transcriptomics may outperform proteomics due to the fact that it has more comprehensive coverage of the genome, and that this includes genes more informative of metastatic mechanisms. We focused specifically on the mean squared error (MSE) metric, as this showed the performance difference between the two modalities. We also proceeded with the linear SVR models for both modalities as described in the main text.

To test our hypothesis, we re-ran our model hyperparameter tuning and fitting pipeline (see Methods for details) for both transcriptomics and proteomics using linear SVRs. However, in this instance, we not only looked at the same sample size (247 samples in common between the two modalities) as in the power analysis comparison (Fig. 2a), we also only used the intersection of features. Due to a lack of 1-to-1 mapping between feature identities across the two modalities, taking the intersection between them results in 10,560 and 10,202 features for proteomics and transcriptomics, respectively.

Examination of feature selection patterns across hyperparameter tuning folds further revealed distinct effective dimensionalities between modalities. Transcriptomic models selected all available features in seven folds, 5,000 features in two folds, and 1,000 features in one fold, whereas proteomic models selected 5,000 features in eight folds, 1,000 features in one fold, and 500 features in one fold. This suggests that predictive signal in transcriptomics is distributed more smoothly across a large number of features, with many genes contributing weak but additive effects. In contrast, proteomic signals appear more concentrated within a smaller subset of features, consistent with lower effective dimensionality. Importantly, although selected features overlapped across modalities (Fig. 2c), when the same feature space was available, overlap was moderate (median Jaccard index = 0.47 across folds). This moderate Jaccard index reflects the asymmetry in the number of features selected by each modality—with transcriptomics often selecting all available features while proteomics selected ~5,000. The Jaccard index, which measures the ratio of shared features to all features selected by either model, is necessarily lower when one model selects substantially more features than the other, even when the smaller set is largely contained within the larger one. This likely reflects variability in the selection of correlated or partially redundant features under regularization—such that different features can serve as interchangeable representatives of similar underlying signals—rather than disagreement in which biological signals are informative. Consistent with this interpretation, strong feature overlap and concordant model coefficients were observed (Fig. 2c-d), indicating agreement in the direction and relative importance of shared features. Together, these results support the view that transcriptomics and proteomics encode largely overlapping predictive information, while differing in the breadth with which that information is distributed across features.

Across the same 10-fold cross-validation train-test splits, we observed that there was no longer a significant difference (paired Wilcoxon signed rank p-value > 0.05) between models fit on the two modalities (Fig. S9a); the effect size of the difference dropped from -1.00 to -0.29. Altogether, these results initially suggested that genome coverage differences might explain transcriptomics' superior performance.

To ensure that this discrepancy was not driven by the specific subset of samples shared between proteomics and transcriptomics, we next asked whether models trained on all available features could recover the power analysis results (Fig. 2a) when evaluated on the same 10-fold cross-validation splits used for the intersecting-feature analysis. In both comparisons, models were fit using all features and an equal sample size; however, in the power analysis, the transcriptomic model was trained on a random subset of all transcriptomic samples, whereas here it was trained exclusively on the subset of samples shared with proteomics (247 of 248 total proteomic samples). In contrast, proteomic models were trained on nearly identical sample sets in both cases (248 versus 247 samples). If performance differences were primarily driven by feature availability, we would expect this comparison to recapitulate the power analysis results. Instead, we again observed no significant difference between modalities (paired Wilcoxon signed-rank test, $p > 0.05$), with an effect size comparable to that observed for the intersecting-feature analysis (Cohen's $d = -0.32$; Fig. S9b).

This prompted us to directly compare model performance between the power analysis and models trained on the proteomics-shared sample subset. We found that transcriptomic models trained on the shared sample subset exhibited virtually identical performance when using all versus intersecting features (Cohen's $d = 0.02$). In contrast, both of these models differed substantially from transcriptomic models trained in the power analysis, with a moderate-to-large effect size (Cohen's $d = -0.77$; Fig. S9c). These shared-sample results were similarly observed for proteomic models (Fig. S9d). Together, these findings indicate that performance differences observed in this dataset (Fig. 2a) are not driven by intrinsic modality-specific effects or feature availability, but instead primarily reflect differences in the samples available to each modality.

### Supplementary Figures

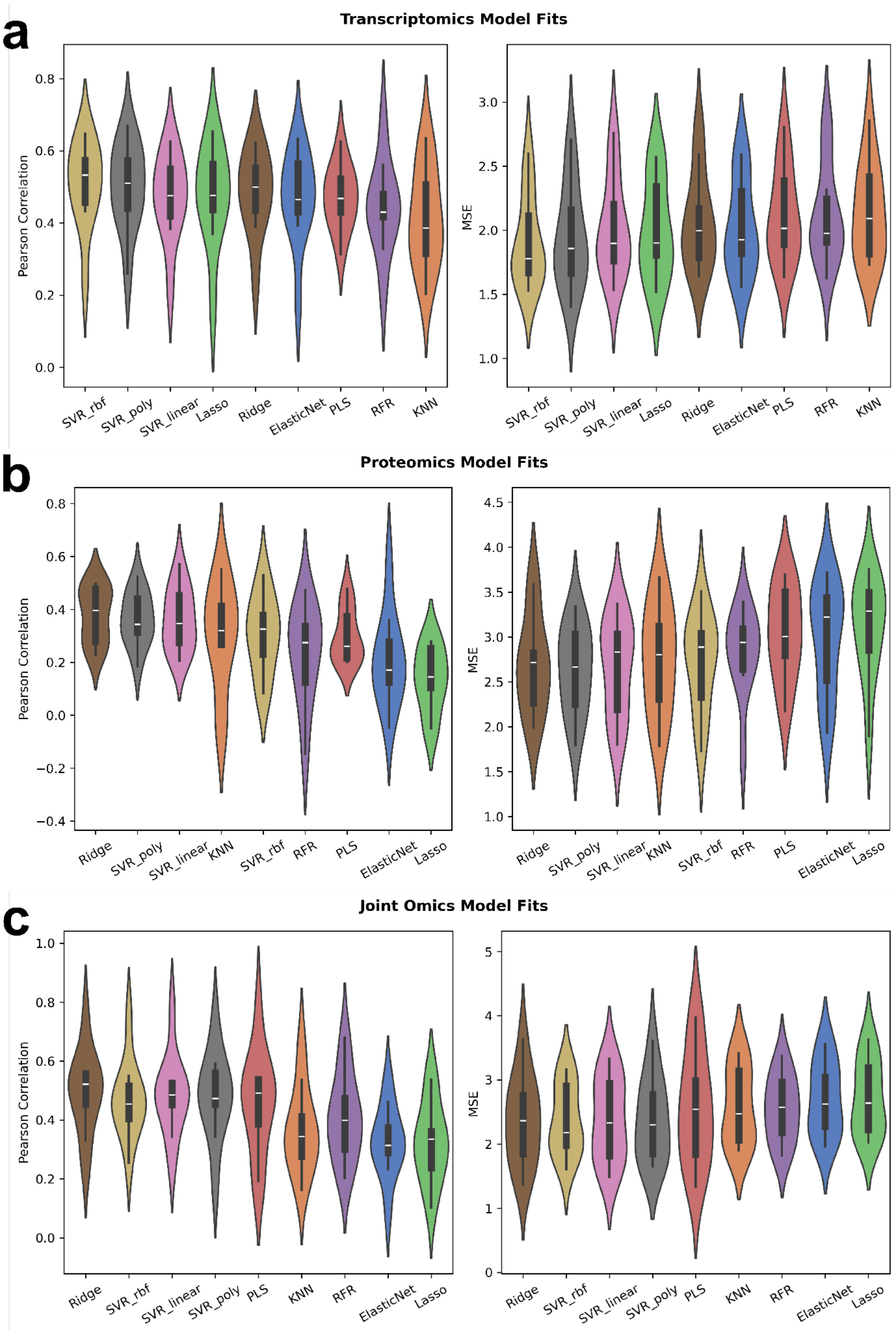


**Fig. S1: Results of 10-fold CV for all hyperparameter tuned models fit on proteomics data.** Violin plots visualize models' predictive performance distribution output from 10-fold CV. Performance is assessed either by Pearson correlation (left panel) or mean squared error (right panel). Performance of 9 predictive models (x-axis) is ordered by consensus ranks of median value. For each fold, specific model hyperparameters were optimized using nested 5-fold CV. All statistical significance is determined using a two-sided Mann-Whitney U (MWU) test and the FDR is controlled for multiple testing using the Benjamini-Hochberg correction (**** $q \leq 10^{-4}$, *** $q \leq 10^{-3}$, ** $q \leq 0.01$, * $q \leq 0.1$). Comparisons that are not significant are not annotated. No comparisons have $q \leq 0.1$. Visualizations are for models fit on **a** transcriptomics, **b** proteomics, and **c** both transcriptomics and proteomics.

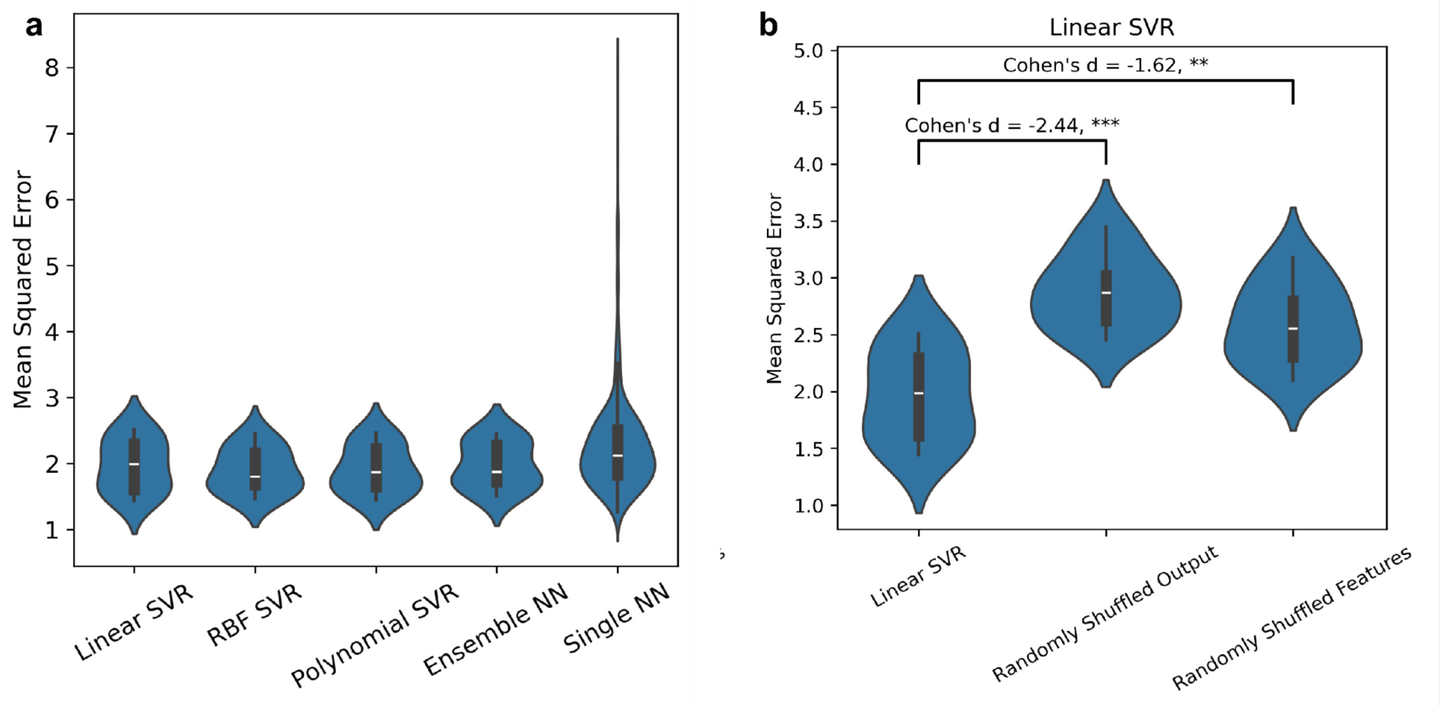


**Fig. S2**: **Transcriptomics model comparisons by MSE**. Violin plots visualize performance distribution output from 10-fold CV. Models' predictive performance is assessed by Mean Squared Error. All statistical significance is determined using a Mann-Whitney U (MWU) test and the FDR is controlled for multiple testing using the Benjamini-Hochberg correction (**** $q \leq 10^{-4}$, *** $q \leq 10^{-3}$, ** $q \leq 0.01$, * $q \leq 0.1$). Comparisons that are not significant are not annotated. **a** Comparison of consensus linear SVR model to other consensus SVR models' and neural nets' (x-axis) performance (y-axis) (two-sided MWU test). No comparisons have $q \leq 0.1$. **b** Consensus Linear SVR model performance (y-axis) compared to random baseline models (x-axis) (one-sided MWU test). This is the MSE analogue to Fig. 1a and 1c.

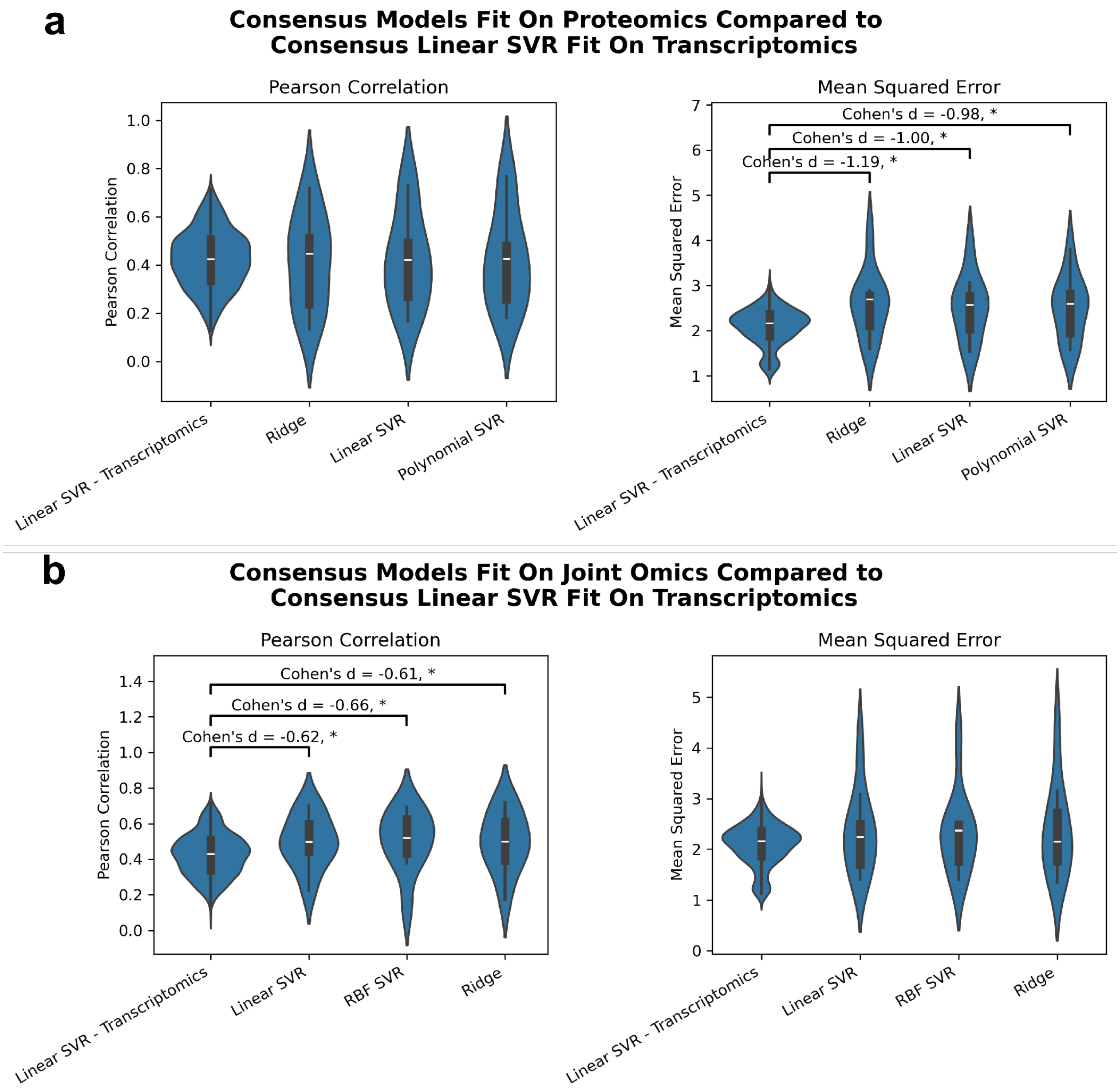


**Fig. S3:** All statistical significance is determined using a Mann-Whitney U (MWU) test and the FDR is controlled for multiple testing using the Benjamini-Hochberg correction (**** $q \le 10^{-4}$, *** $q \le 10^{-3}$, ** $q \le 0.01$, * $q \le 0.1$). Comparisons that are not significant are not annotated. Models' predictive performance is assessed either by Pearson correlation (left panels) or mean squared error (right panels). Violin plots visualize performance distribution output from 10-fold CV. Violin Plots of the performance of consensus linear SVR model at the same sample size as **a** proteomics data (248 samples) or **b** joint omics data (247 samples) assessed by power analysis and top-performing consensus proteomics models. These Violin Plots illustrate in more detail the same information visualized in Fig. 2a.

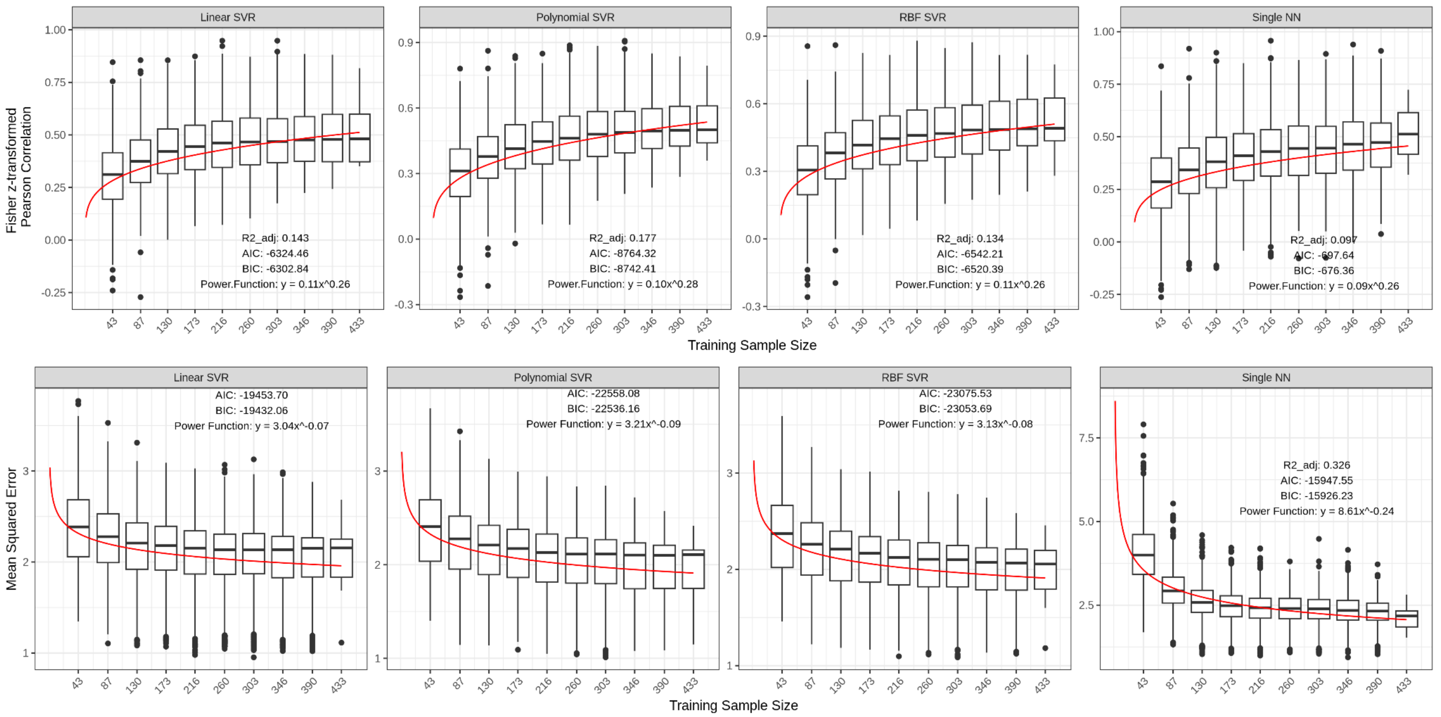


**Fig. S4: Results of power analysis on consensus transcriptomic models.** Box Plots of individual consensus models fit on transcriptomics data. Model performance (y-axis) assessed by Fisher z-transformed Pearson correlation (top panels) or mean squared error (bottom panels) across training sample size (x-axis). Red lines visualize the power function regression fits for each model type and performance metric. Plots are annotated with assessment metrics for these regression fits as well as regression parameters.

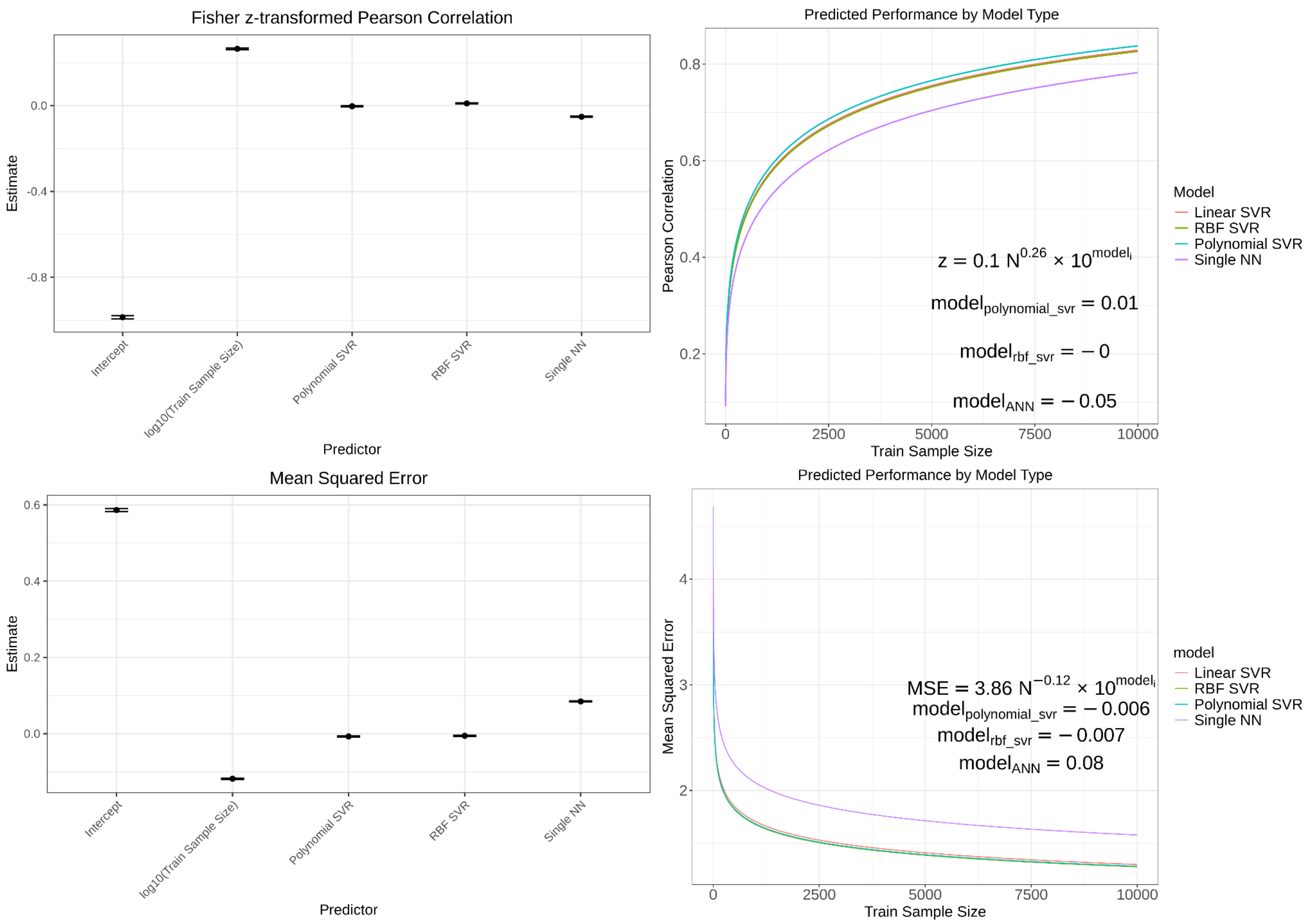


**Fig. S5: Power function fits for performance of consensus transcriptomic models across sample size.** Performance of consensus models by Pearson correlation (top panels) and MSE (bottom panels) estimated by power function regression fits that include the model type as a covariate. Left panels show estimates (y-axis, scatter point) and standard errors (horizontal bars) for each parameter (x-axis) in the model prior to back transformation (exponentiation and, in the case of Pearson correlation, inverse Fisher z-transformation). With the exception of the RBF SVR for Pearson, all estimates are significant (two-sided Student's t-test, p-value ≤ 4.12e-5); the non-significant estimate indicates no detectable difference from the reference linear SVR. In a power function $y = ax^b$, the intercept represents $\log_{10}(a)$, "log10(Train Sample Size)" represents $b$, and the remaining estimates represent the effect of the model type with respect to the Linear SVR as a baseline (see Methods for details). Right panels visualize predicted performance (y-axis) for each model type across training sample size (x-axis) from 1 to 10,000 training samples. Plots are annotated with the back-transformed power function fitted parameters, with $N$ representing training sample size and $model_i$ representing the effect of the model type with respect to Linear SVR ($model_{linear\_svr} = 0$, $10^0 = 1$, no effect). The correlation function annotation is in terms of the Fisher z-transformed Pearson correlation ($z$), whereas the curves and y-axis are back-transformed to Pearson correlation.

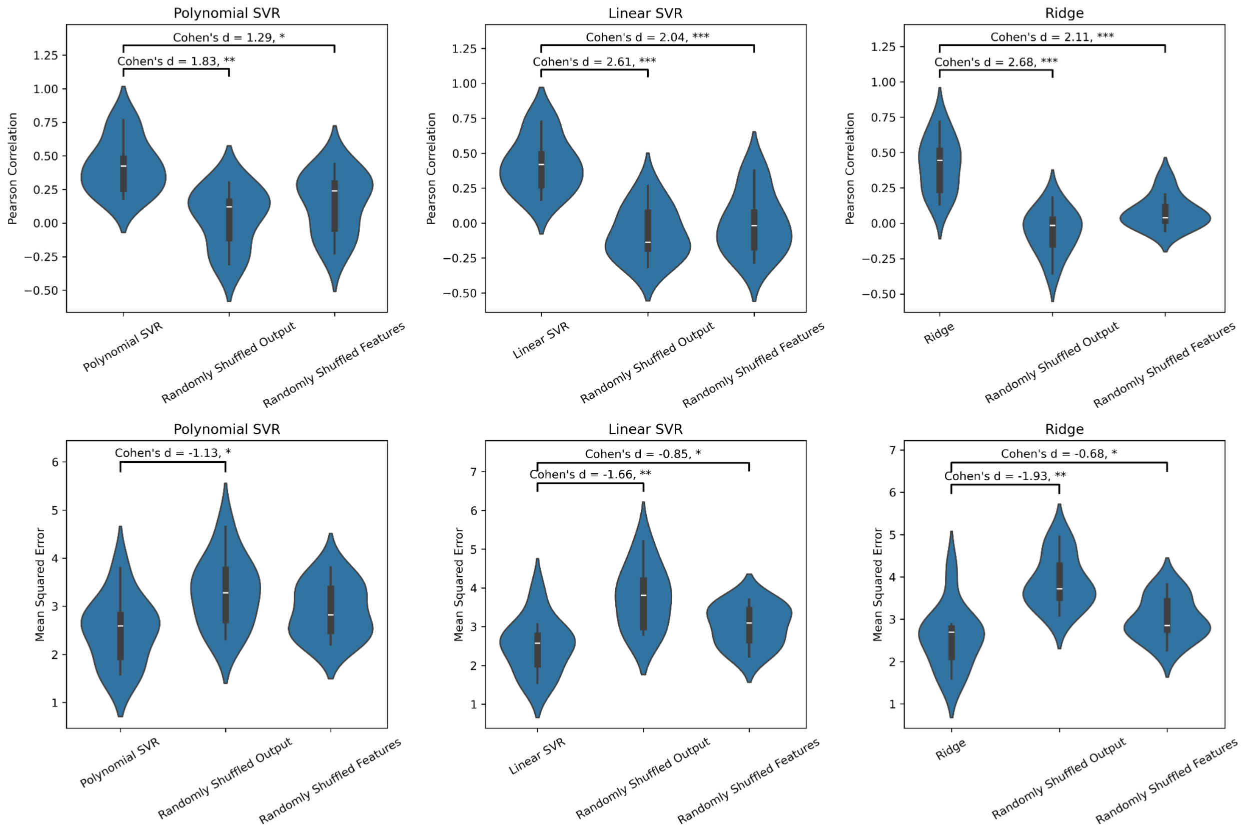


**Fig. S6: Comparison of consensus proteomics models to random baselines**. All statistical significance is determined using a Mann-Whitney U (MWU) test and the FDR is controlled for multiple testing using the Benjamini-Hochberg correction (**** $q \leq 10^{-4}$, *** $q \leq 10^{-3}$, ** $q \leq 0.01$, * $q \leq 0.1$). Comparisons that are not significant are not annotated. Statistical tests are one-sided (testing whether the actual model performs better than random) unless otherwise specified. Models' predictive performance is assessed either by Pearson correlation (top panels) or mean squared error (bottom panels). Violin Plots display performance distributions output from 10-fold CV.

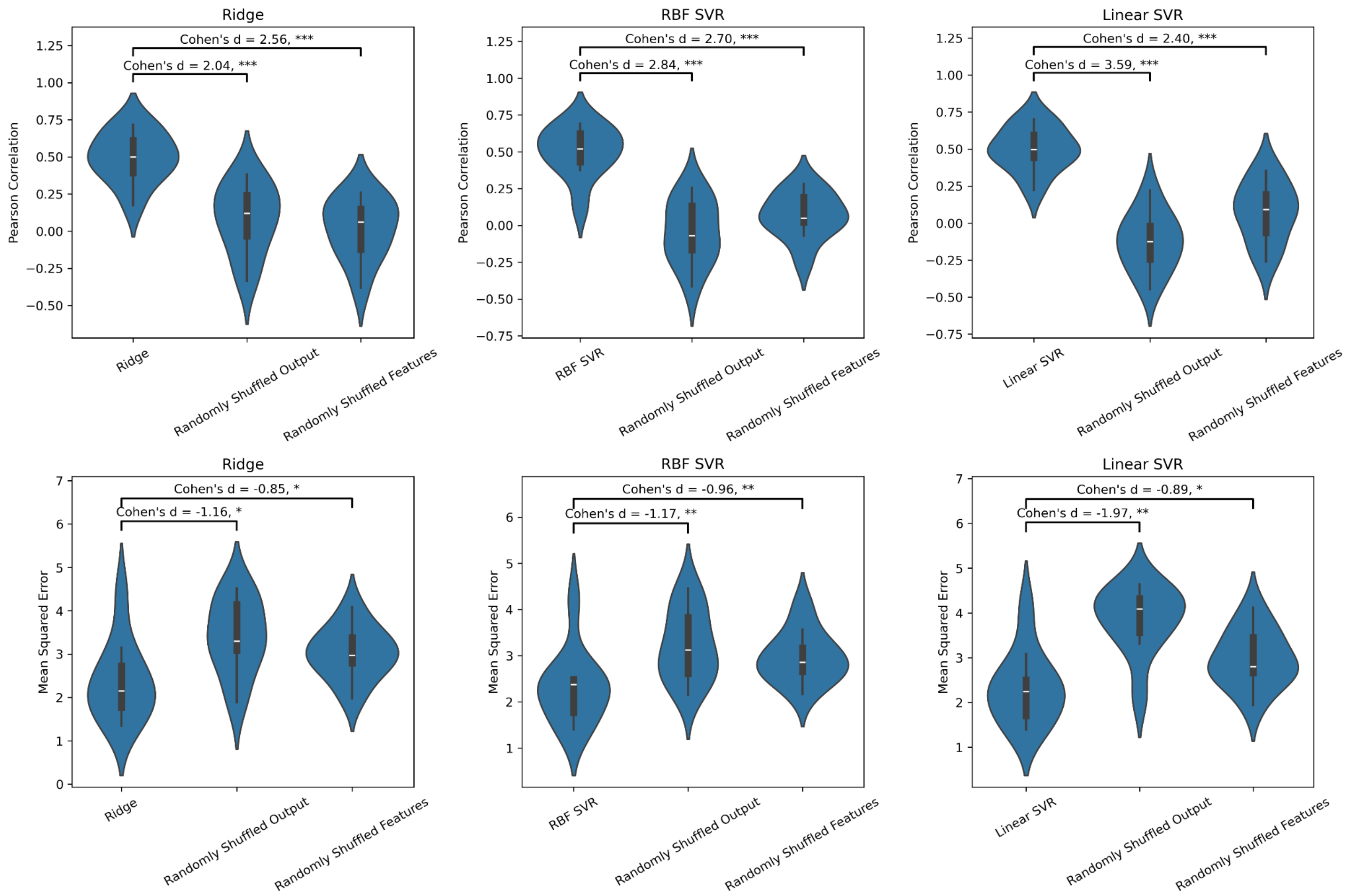


**Fig. S7: Comparison of consensus joint omics models to random baselines**. All statistical significance is determined using a Mann-Whitney U (MWU) test and the FDR is controlled for multiple testing using the Benjamini-Hochberg correction (**** $q \leq 10^{-4}$, *** $q \leq 10^{-3}$, ** $q \leq 0.01$, * $q \leq 0.1$). Comparisons that are not significant are not annotated. Statistical tests are one-sided (testing whether the actual model performs better than random). Models' predictive performance is assessed either by Pearson correlation (top panels) or mean squared error (bottom panels). Violin Plots display performance distributions output from 10-fold CV.

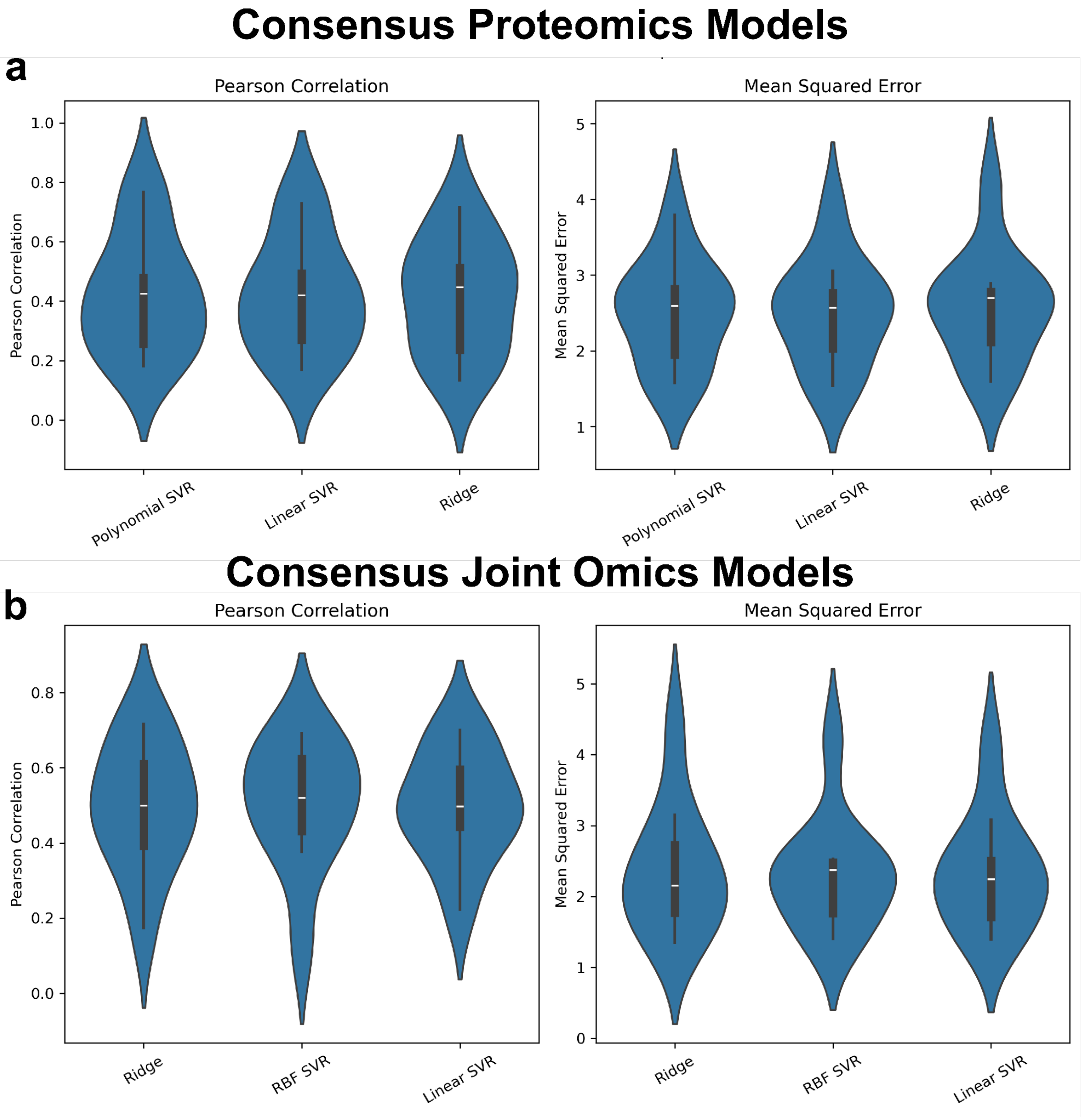


**Fig. S8: Comparison of performance between consensus models**. All statistical significance is determined using a Mann-Whitney U (MWU) test and the FDR is controlled for multiple testing using the Benjamini-Hochberg correction (**** $q \leq 10^{-4}$, *** $q \leq 10^{-3}$, ** $q \leq 0.01$, * $q \leq 0.1$). Comparisons that are not significant are not annotated. Statistical tests are two-sided. Models' predictive performance is assessed either by Pearson correlation (left panel) or mean squared error (right panel). Violin Plots display performance distributions output from 10-fold CV. This is analogous to Fig. 1a in the main text for **a** proteomics and **b** transcriptomic and proteomics combined.

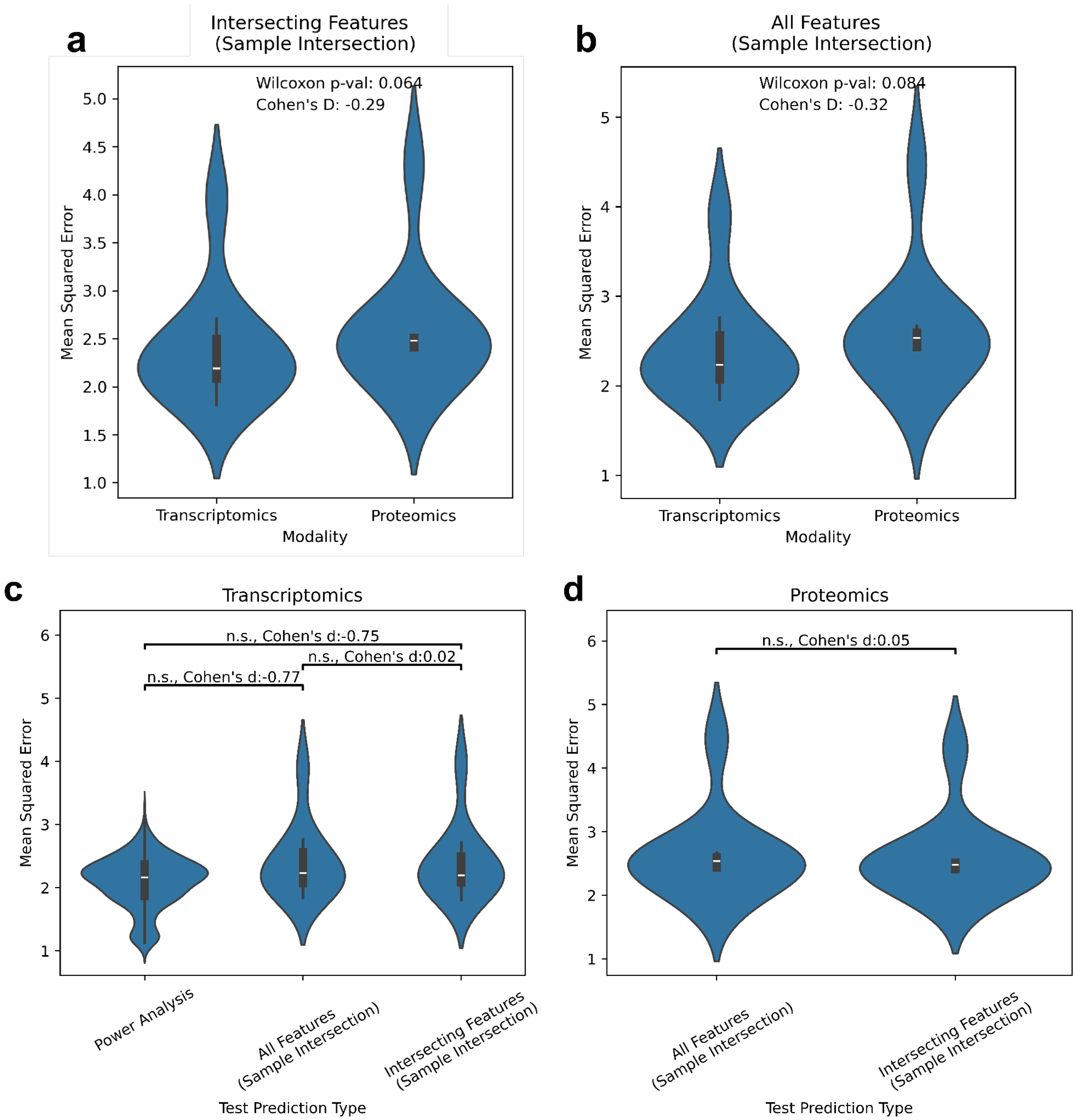


**Fig. S9: Further comparison of transcriptomics and proteomics model fits.** Violin plots visualize performance distribution from 10-fold CV measured by MSE (y-axis) comparing linear SVR performance fit on transcriptomics or proteomics (x-axis) **a** Models fit on intersecting features and samples between both modalities. The hyperparameter tuning and subsequent 10-fold CV pipeline was run as described for all other consensus models. **b** Analogous comparison as in (a), including the same 10-fold split, but using hyperparameters identified for consensus models fit on all features and samples as in the main text. **c** Comparison of transcriptomic model performances from (a) and (c) to that from the power analysis on the same sample size. **d** Comparison of proteomic model performances from (a) and (b).

### References

1. Lakshminarayanan, B., Pritzel, A. & Blundell, C. Simple and Scalable Predictive Uncertainty Estimation using Deep Ensembles. in *Advances in Neural Information Processing Systems* 6402–6413.
